# Cell lineage-dependent chiral actomyosin flows drive cellular rearrangements in early development

**DOI:** 10.1101/842922

**Authors:** Lokesh Pimpale, Teije C. Middelkoop, Alexander Mietke, Stephan W. Grill

**Affiliations:** Max Planck Institute of Molecular Cell Biology and Genetics, Dresden, Germany; Biotechnology Center, TU Dresden, Dresden, Germany; Excellence Cluster Physics of Life, TU Dresden, Germany; Max Planck Institute for the Physics of Complex Systems, 01187 Dresden, Germany; Chair of Scientific Computing for Systems Biology, Faculty of Computer Science, TU Dresden, 01187 Dresden, Germany; Center for Systems Biology Dresden, 01307 Dresden, Germany; Department of Mathematics, Massachusetts Institute of Technology, 77 Massachusetts Avenue, Cambridge, Massachusetts 02139-4307, USA

## Abstract

Proper positioning of cells is important for many aspects of embryonic development, tissue homeostasis, and regeneration. A simple mechanism by which cell positions can be specified is via orienting the cell division axis. This axis is specified at the onset of cytokinesis, but can be reoriented as cytokinesis proceeds. Rotatory actomyosin flows have been implied in specifying and reorienting the cell division axis in certain cases, but how general such reorientation events are, and how they are controlled, remains unclear. In this study, we set out to address these questions by investigating early *Caenorhabditis elegans* development. In particular, we determined which of the early embryonic cell divisions exhibit chiral counter-rotating actomyosin flows, and which do not. We follow the first nine divisions of the early embryo, and discover that chiral counter-rotating flows arise systematically in the early AB lineage, but not in early P/EMS lineage cell divisions. Combining our experiments with thin film active chiral fluid theory we identify specific properties of the actomyosin cortex in the symmetric AB lineage divisions that favor chiral counter-rotating actomyosin flows of the two halves of the dividing cell. Finally, we show that these counter-rotations are the driving force of both the AB lineage spindle skew and cell reorientation events. In conclusion, we here have shed light on the physical basis of lineage-specific actomyosin-based processes that drive chiral morphogenesis during development.

## INTRODUCTION

Proper positioning of cells is instrumental for many aspects of early embryonic development [1–3]. One way to achieve proper cell positioning is via migration [4, 5]. Another way by which cell positions can be specified is via the orientation of the cell division axis [6–8]. Since the orientation of the mitotic spindle dictates the cell division axis, the orientation of the mitotic spindle at cleavage is instrumental for determining daughter cell positions [9–12]. This axis can be set at the beginning of cytokinesis, by assembling the mitotic spindle in the correct orientation from the start, or during cytokinesis by reorientation of the mitotic spindle [13–18] While several studies highlight the mechanisms that determine the initial orientation of the mitotic spindle at the onset of cytokinesis [15–17, 19, 20] the mechanisms controlling reorientation remain not very well understood. We here set out to investigate cell repositioning during cytokinesis in the early development of the *C. elegans* nematode. The fully grown hermaphrodite worm consists of exactly 959 somatic cells that are essentially invariant both in terms of position and lineage [21–24]. Development is deterministic from the start: the one-cell embryo undergoes an asymmetric cell division that gives rise to the AB (somatic) lineage and the P lineage [22, 25]. While the anterior daughter cell, AB, undergoes a symmetric cell division into ABa and ABp, the posterior daughter cell, P_1_, divides asymmetrically into EMS forming the endoderm and mesoderm, and P_2_ forming the germ line [22]. Appropriate cell-cell contacts are instrumental for development as they can determine cell identity [26–30]. For example, reorientation of the ABa and ABp cells via pushing with a micro needle leads to an altered cell-cell contact pattern and an altered body plan with an inverted L/R body axis [31]. Consequentially, proper cell positioning, perhaps mediated via repositioning of the mitotic spindle during cytokinesis, is crucial [32]. Here, we set out to investigate which of the cells of the early embryo undergo reorientations during cytokinesis, and by which mechanism they do so.

Recently, a role for the actomyosin cell cortex in determining the cell division axis of early *C. elegans* blastomeres was identified [17, 33]. The actomyosin cortex is a thin layer below the plasma membrane that consists mainly of actin filaments, actin binding proteins and myosin motor proteins [34]. Collectively, these molecules generate contractile forces that can shape the cell, drive cortical flows during polarization and orchestrate other active processes such as cell division [35, 36]. Cell-cell contacts can impact on the activity of myosin and the generation of contractile stresses, and the resultant pattern of cortical flows can determine the orientation of the mitotic spindle at the onset of cytokinesis [17]. From a physical point of view, the actomyosin cortex can be thought of as a thin layer of a mechanically active fluid [35, 37– 39] with myosin-driven active stress gradients generating cortical flows [33, 35]. Interestingly, actomyosin can also exhibit rotatory flows driven by active torque generation [40, 41]. These chiral rotatory cortical flows reorient the ABa cell and the ABp cell during cytokinesis, driving a cell skew of ∼ 20° during division [33]. This skew results in a L/R asymmetric cell-cell contact pattern [42], thus executing left-right (L/R) symmetry breaking in the entire organism. However, how general such reorientation events are, and how they are controlled, remains unclear. Furthermore, whether chiral flows are prevalent in other cell divisions as well, and if they play a prominent role in cell repositioning during early embryogenesis of the *C. elegans* nematode, remains poorly understood.

## RESULTS

We set out to determine the extent of chiral rotatory movements in the actomyosin cell cortex of the first nine cell division in early *C. elegans* development. We started with investigating the first two divisions, the divisions of the P_0_ zygote and the AB cell, and quantified flows from embryos containing endogenously tagged non-muscle myosin-II (NMY-2)::GFP using spinning disc microscopy (Fig. 1(a), Supplement Video 1,2). We used particle image velocimetry (PIV) to determine cortical flow velocities in two rectangular ROI’s placed on opposite sides of the ingressing cytokinetic furrow (Fig. 1(a)). The velocity component parallel (y-direction) to the plane of cytokinesis (as determined by the cytokinetic ring) was averaged over the area of each box and over a time period of 21s after the onset of furrow ingression, indicating the beginning of cytokinesis. Strikingly, while P_0_ does not exhibit counter-rotating flows, AB exhibits counterrotating flows with the two dividing halves spinning in opposite directions during division (Supplement Video 1,2). To quantify the speed and the handedness of these rotary flows, we use the velocities measured in each box (<u>v_1_</u> and <u>v_2_</u>) to define a chiral counter-rotation flow velocity v_c_ = <u>e_z_</u> · (<u>e_x_</u> × <u>v_2_</u> − <u>e_x_</u> × <u>v_1_</u>) (akin to [33]), where <u>e_x_</u> is a unit length base vector pointing from the cytokinetic ring towards the pole, e_y_ is an orthogonal unit length vector parallel to the cytokinetic plane (Fig. 1(a)), and <u>e_z_</u> = <u>e_x_</u> × <u>e_y_</u>. With this definition, |v_c_| quantifies the speed of counter-rotating flows and the sign of v_c_ denotes their handedness (see methods and Supplement Fig. 1(a)). Indeed, counter-rotating flows are absent during the division of the P_0_ cell with v_c_ = 0.01 ± 0.51 *µ*m/min (mean ± error of the mean at 95% confidence unless otherwise noted), but present during AB cell division with v_c_ = −7.01 ± 0.34 *µ*m/min. We conclude that of first two cell divisions in the developing nematode, one displays counter-rotating actomyosin flows during cytokinesis while one does not.

**Figure 1:**
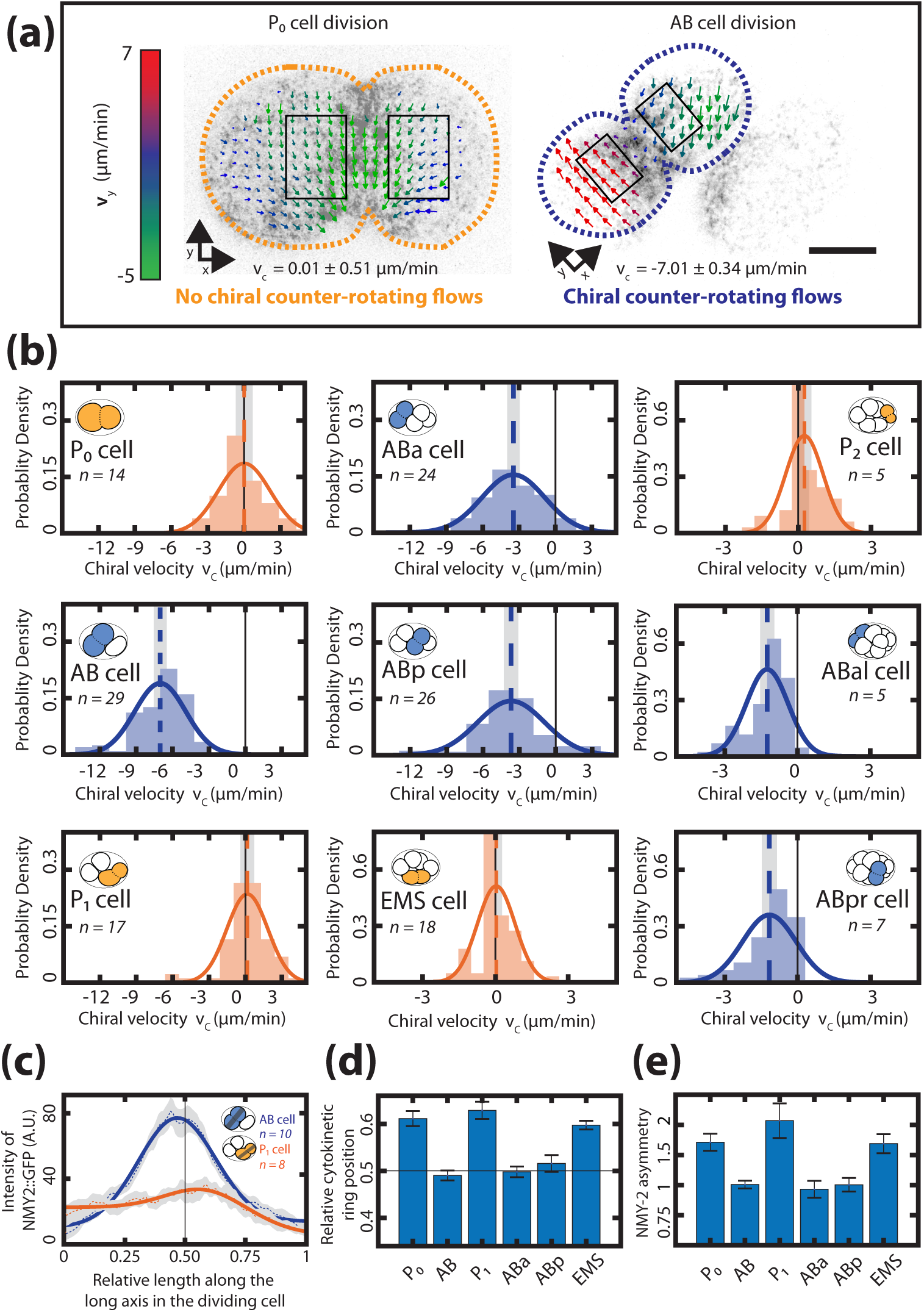
Chiral counter-rotating flows are cell-lineage specific. **(a)**, Representative images of cortical myosin (grey, NMY-2::GFP) for the first (P_0_, yellow) and the second (AB, blue) cell division of *C. elegans* embryos. Arrows indicate the cortical flow field as measured by PIV and time-averaged over 21 s and over the onset of cytokinesis. Arrow colors indicate the y-direction velocity (parallel to the cytokinesis furrow, coordinate systems are indicated). Dotted lines indicate cell boundaries, black boxes represent the regions of interest used for calculating velocities. Scale bar, 10 *µ*m. **(b)**, Histograms of the instantaneous chiral counter-rotating flow velocity v_c_ (see methods for definition). Solid lines indicate the best-fit gaussian probability density function. Dotted vertical colored lines represent the mean v_c_, grey boxes represent the error of the mean. Thin black solid lines indicate a chiral flow velocity of zero. Inset, colored cells indicate the cell analyzed; AB lineage in blue, P/EMS lineage in yellow. **(c)**, Dotted lines with shaded error region indicate the average myosin concentration profile along the long axis of the AB (blue) and P_1_ (orange) dividing cell, solid lines represent a best fit with a combined step and gaussian function (see methods) Inset, colored cells indicate the cell analyzed; grey stripe indicates the region used for averaging. **(d)**, Average relative position of the cytokinetic ring along the long axis of the dividing cell for the first 6 cell divisions. Black thin lines in **(c)** and **(d)** indicate the center of the cell.**(e)**, Myosin asymmetry ratio (Anterior [NMY-2::GFP] / Posterior [NMY-2::GFP]) (see methods) for the first 6 cell divisions along the long axis of the dividing cell. Errors indicate the error of the mean at 95% confidence.

This raises the question which of the later cell divisions display chiral counter-rotating flows, and which do not. To this end, we quantified chiral counter-rotating flows for the next seven cell divisions as described above. We find that cells of the AB lineage (ABa, ABp, ABar and ABpr) counter-rotate with average velocities of v_c_ = −3.46 ± 0.33 *µ*m/min, −3.68 ± 0.41 *µ*m/min, −1.24 ± 0.19 *µ*m/min and −1.16 ± 0.22 *µ*m/min respectively (Fig. 1(b)). In contrast, average counter-rotating flow velocities of cells of the P/EMS lineage (P_1_, EMS and P_2_) are 0.15 ± 0.38 *µ*m/min, 0.03 ± 0.14 *µ*m/min and 0.253 ± 0.19 *µ*m/min, respectively (Fig. 1(b)). We also observe that cells of the P/EMS lineage, but not those of the AB lineage, exhibit a whole-cell (net-)rotating flow where the cortex in both cell halves move in the same y-direction [43] (Supplement Fig. 2(b), Supplement Video 1-5). To conclude, early cell divisions of the AB lineage, but not of the P/EMS lineage, undergo chiral counter-rotating flows. Note also that the chiral counter-rotating flow velocity in the AB lineage decreases as development progresses and cells become smaller (Fig. 2(b)). Together these findings show that both presence and strength of chiral counter-rotating flows are related to the fate of the dividing cell.

**Figure 2:**
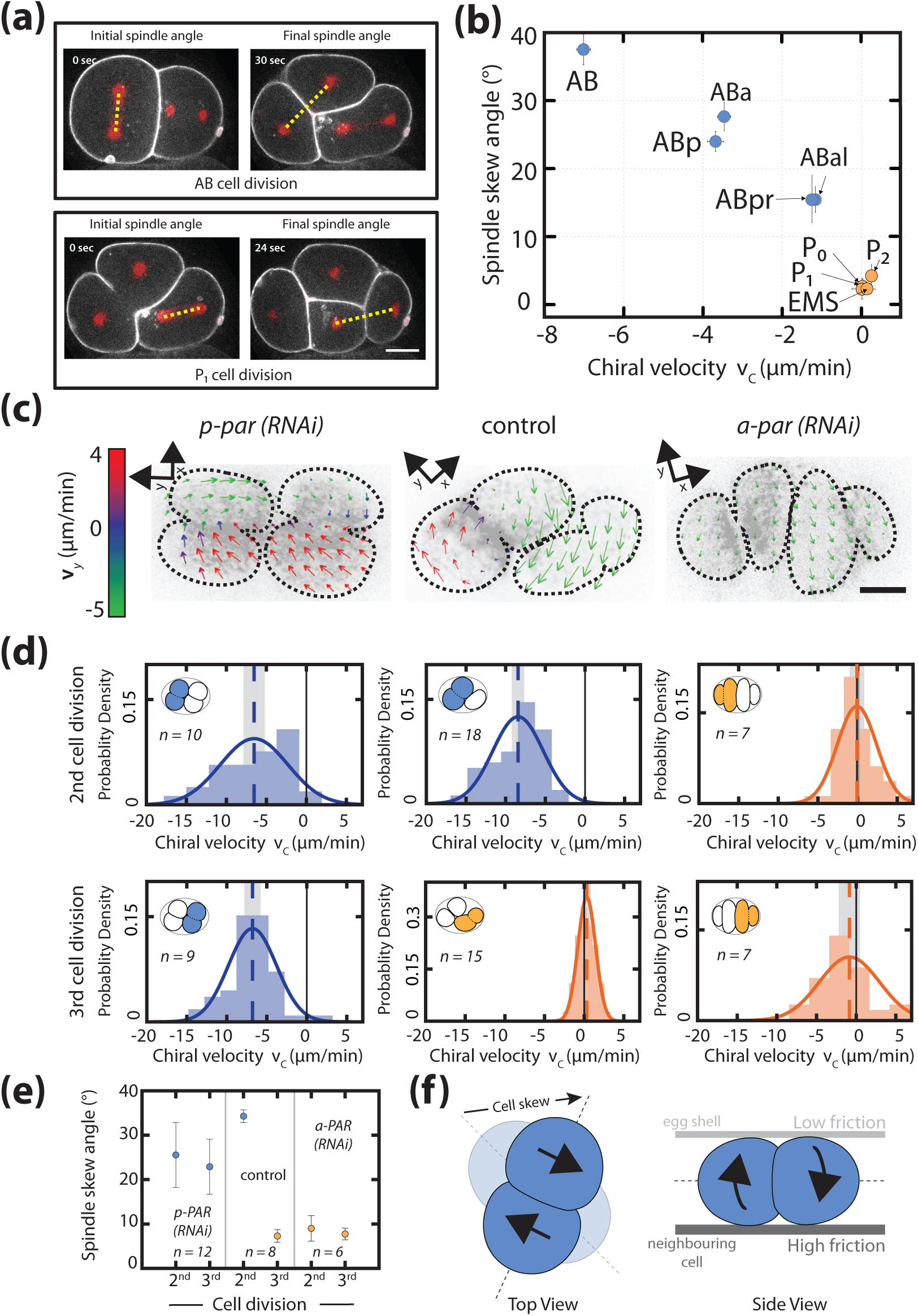
Chiral counter-rotating flows are cell-lineage dependent. **(a)**, Representative images show the angle of the mitotic spindle (yellow dotted line) before the onset (left) and at the end (right) of the second (AB, top) and the third (P_1_, bottom) cell division. White, cell membrane imaged by PH::GFP; red, spindle poles as imaged by TBB-2::mCherry. **(b)**, Spindle skew angle (final minus initial spindle angle) vs. chiral velocity for the first nine cell divisions (N = 8 for all cell diviisons). **(c)**, Representative images of cortical myosin (grey, NMY-2::GFP) for the second and the third cell division of *C. elegans* embryos. Arrows indicate the cortical flow field as measured by PIV and time-averaged over 21 s and over the onset of cytokinesis. Arrow colors indicate the y-direction velocity (parallel to the cytokinesis furrow, coordinate systems are indicated). **(d)**, Histograms of the instantaneous chiral counter-rotating flow velocity v_c_ for *par* perturbations (see methods for definition). Solid lines indicate the best-fit gaussian probability density function. Dotted vertical colored lines represent the mean v_c_, grey boxes represent the error of the mean. Thin black solid lines indicate a chiral flow velocity of zero. Inset, colored cells indicate the cell analyzed; AB lineage in blue, P/EMS lineage in orange. **(e)**, Spindle skew angle for control (L4440), *par-2; chin-1; lgl-1(RNAi)* and *par-6(RNAi)* embryos. **(f)**, Sketch of a dividing cell undergoing cell skew via chiral counter-rotating flows, left: top view, right: side view. Chiral counter-rotation of the two halves of a dividing cell gives rise to a cell skew if the dividing cell experiences non-uniform friction with the other surfaces that it is in contact with, e.g. low friction with an eggshell above and high friction with a cell below. Scale bars, 10 *µ*m. Errors indicate the error of the mean at 95% confidence.

We have shown previously that chiral rotating cortical flows are driven by active torque generation that depends on the myosin distribution [33]. Given that chiral counter-rotating flows arise only during AB and not P/EMS cell divisions, we hypothesize that myosin distributions during division of the AB and the P/EMS-lineage cells differ in specific features. To investigate this, we determined the NMY-2 distribution along the long axis of dividing cells (Fig. 1(c)) for the first six cell divisions, and extracted two features: a myosin ratio that characterizes the difference in myosin intensity between the two cell halves and the relative cytokinetic ring position that characterizes the relative position of the myosin peak along the long axis, and the degree of asymmetry of cell division. We find that myosin ratios for the AB, ABa and Abp cells are 1.0 ± 0.13, 0.92 ± 0.29, and 1.00 ± 0.24, respectively, indicating that there is no significant difference in cortical myosin concentration between the two halves of AP lineage dividing cells. In contrast, the myosin ratios in P_0_, P_1_ and EMS are 1.67 ± 0.3, 2.07 ± 0.61, 1.68 ± 0.33, respectively (Fig. 1(e)). We find that cytokinetic ring position for AB, ABa and ABp cell are at positions of 0.49 ± 0.01, 0.49 ± 0.01 and 0.51 ± 0.02 relative to cell length, respectively. In contrast, the cytokinetic ring position for P_0_, P_1_ and EMS cells are at positions of 0.58 ± 0.01, 0.57 ± 0.02 and 0.59 ± 0.01 relative to cell length, respectively (Fig. 1(d)). To conclude, the early cell divisions of the AB lineage are symmetric in terms of myosin intensity ratio and ring position, while early cell divisions of the P/EMS lineage are asymmetric in these two features.

We next asked if the differences in presence and absence of chiral counter-rotating flows between AB and P/EMS cells can be accounted for by the observed difference in myosin ratio and cytokinetic ring position. We utilized thin film active chiral fluid theory [33, 44] in an isotropic model cell of similar dimension with constant coefficient of friction with respect to the cytoplasm of the cell and the outside environment, and determined the chiral counter-rotating flow velocity v_c_ for different values of myosin ratio and ring position. We find that a myosin ratio of one and a symmetric ring position at 0.5 results in a maximal v_c_, while myosin ratios different from one and asymmetric ring position results in lower values of v_c_ (Supplement Fig. 3(d)-(e)). This indicates that the switch between chiral and non-chiral flows between the two lineages could in principle be attributed to the observed difference in myosin ratio, cytokinetic ring position or both. We note, however, that while thin film active chiral fluid theory can relate at a quantitative level the myosin profile along the long axis of the dividing cell with the chiral y-component of the actomyosin flow-field for AB, ABa, and ABp, this is not possible for the P/EMS lineage (Supplement Fig. 3(a)-(b), Supplement Table 1,2). We suspect this is due to specific features of the overall dynamics that are not included in our theoretical description, such as movements of the midbody remnant particular to the P/EMS linage [45] or possible inhomogeneities in the friction coefficient (e.g. friction with respect to the neighboring cells). This indicates that our understanding of the mechanisms that shape the cortical flow fields in these divisions is still incomplete.

We next focus our attention on the mechanism by which dividing cells assume their correct position inside the developing embryo. Specific rotations of the spindle (termed spindle skews) are thought to be a key mechanism by which dividing cells reposition themselves in the embryo [33, 45, 46]. One hypothesis is that spindle skews are driven by spindle elongation [32]. In this case spindle elongation causes a skew because the spindle rearranges within an asymmetric cell shape, thus responding to the constraints provided by neighboring cells and the eggshell (Supplement Fig. 4(a)). An alternative hypothesis is that spindle skews are driven by counter-rotating flows of the dividing cell halves [33]. In this case the mechanism is akin to a bulldozer rotating on the spot by spinning its chains in opposite directions [33]. Counter-rotating cortical flows of the two cell halves will lead to a torque and a cell skew in the case that the cell experiences different friction coefficients on different surfaces (e.g. friction with respect to eggshell in comparison with friction with respect to a neighboring cell; Fig. 2(f)). To discriminate between these two hypothesis, we first asked if those cells that undergo spindle skews also exhibit chiral counter-rotating flows. We quantified spindle skews by determining the positions of spindle poles at the beginning and at the end of cytokinesis in embryos expressing TBB-2::mCherry for the first 11 cell divisions (Supplement Video 9). The angle between the lines that join the two spindle poles at the onset and after cell division defines the spindle skew angle (Fig. 2(a)). We find that cells of the early AB lineage undergo an average skew angle of 21.17 ± 3.22° (AB: 37.509 ± 2.51°; ABa: 27.64 ± 2.29°, ABp: 23.98°, ABal: 17.73± 1.98°, ABar: 15.51 ± 3.51°, ABpl: 10.43 ± 1.92° and ABpr:15.42 ± 2.11°). In comparison, P/EMS lineage cells undergo significantly reduced spindle skews with an average skew angles of of 2.32 ± 0.43° (P_0_: 2.31 ± 1.67°, P_1_: 2.25 ± 0.66°, EMS: 2.66 ± 1.25°, P_2_: 4.36 ± 1.67°). Note also that spindle skew angles in the AB lineage decrease as development progresses, while the P/EMS lineage spindle skew angles remain small and approximately constant (Fig. 2(b)). Given that AB lineage cells, but not P/EMS lineage cells, undergo chiral counter-rotating flows, we conclude that cells that exhibit significant chiral counter-rotating flows also undergo significant spindle skews during early development (Fig. 2(b)).

The results obtained so far are consistent with a scenario in which cell fate determines the presence of chiral counter-rotating flows, and these counter-rotating flows then drive spindle skews. We test for these two causal relationships separately. We first asked if cell fate determines the presence of chiral counter-rotating flows. In this case, we expect that on the one hand, anteriorizing the worm leads to equal and significant counter-rotating flows for the second and third division. On the other hand, we expect that posteriorizing the worm would lead to absence of chiral counter-rotating flows. Consistent with the first expectation, anteriorizing the embryo via *par-2; chin-1; lgl-1 (RNAi)* [47] results in the second and the third cell division exhibiting chiral counter-rotating velocity with v_c_ values that are not significantly different from one another (−6.53 ± 1.13 *µ*m/min and −6.67 ± 0.8 *µ*m/min, respectively; Supplement Video 6; Fig. 2(c)-(d)). Consistent with the second expectation, posteriorizing the embryo via *par-6 (RNAi)* [48] results in both the second and the third cell division not exhibiting counter-rotating flows (v_c_ = 0.07 ± 0.71 *µ*m/min and −0.85 ± 1.13 *µ*m/min, respectively; Supplement Video 8; Fig. 2(c)-(d)). Hence, cell fate determines the presence or absence of chiral counter-rotating flows. Note also that switching the cell fate results in a concomitant changes in the spindle skew angle for both the second and third cell division (Fig. 2(e)). We conclude that cell fate determines both the presence of chiral counter-rotating flows and the degree of spindle skew.

We next tested the second causal relationship, and asked if chiral counter-rotating flows in the cortex drive spindle skews. Changing the speed at which the chains of a bulldozer counter-rotate leads to a change in the speed at which the entire bulldozer rotates. Hence, we evaluated if increasing or decreasing the chiral counter-rotating flow velocity results in increased or reduced rates of spindle skews. We first evaluated if we can increase and decrease counter-rotating flow velocity in the AB cell, by RNAi of RhoA regulators [33]. Weak perturbation RNAi of *ect-2* results in a counter-rotating flow velocity of −3.57 ± 0.57 *µ*m/min, which is reduced in comparison to the unperturbed embryo (v_c_ = −6.6 ± 0.37 *µ*m/min; Fig. 3(a)-(b)). Conversely, weak perturbation RNAi of *rga-3* increases the chiral counter-rotating flow velocity to v_c_ = −8.71 ± 0.58 *µ*m/min (Fig. 3(a)-(b)). Note that these perturbations impact only on active torque generation and chiral counter-rotating flows, and do not significantly change on-axis contractility-driven flow consistent with previous findings [33] (Fig. 3(c)). We now use these perturbations to test whether chiral counter-rotating flows determine the rate of spindle skew. We indeed find that decreasing the chiral counter-rotating flow velocity by mild *ect-2 (RNAi)* leads to concomitant reduction of the rate of spindle skew (average peak rate of spindle skew: 0.59 ± 0.05 °/min (Supplement Video 10) as compared to 1.01 ± 0.1 °/min in control embryos (Supplement Video 11); Fig. 3(d)). Conversely increasing RhoA signaling by mild *rga-3 (RNAi)* treatment leads to increase of both the chiral counter-rotating flow velocity and the rate of spindle skew (average peak rate of spindle skew: 1.29 ± 0.12 °/min (Supplement Video 12); Fig. 3(d)). Note that the peak rates of spindle elongation remain unchanged as compared to the control for both perturbations (Fig. 3(d)). We conclude that changing counter-rotating flow velocities results in concomitant changes of the rates of spindle skews without impacting on the dynamics of spindle elongation. Hence, it is unlikely that spindle skews are driven by the spindle elongating in an asymmetric cell shape [13, 32], as stated in the first hypothesis above. These results instead lead credence to the second hypothesis, that chiral counter-rotating flows mechanically drive cell and spindle skews.

**Figure 3:**
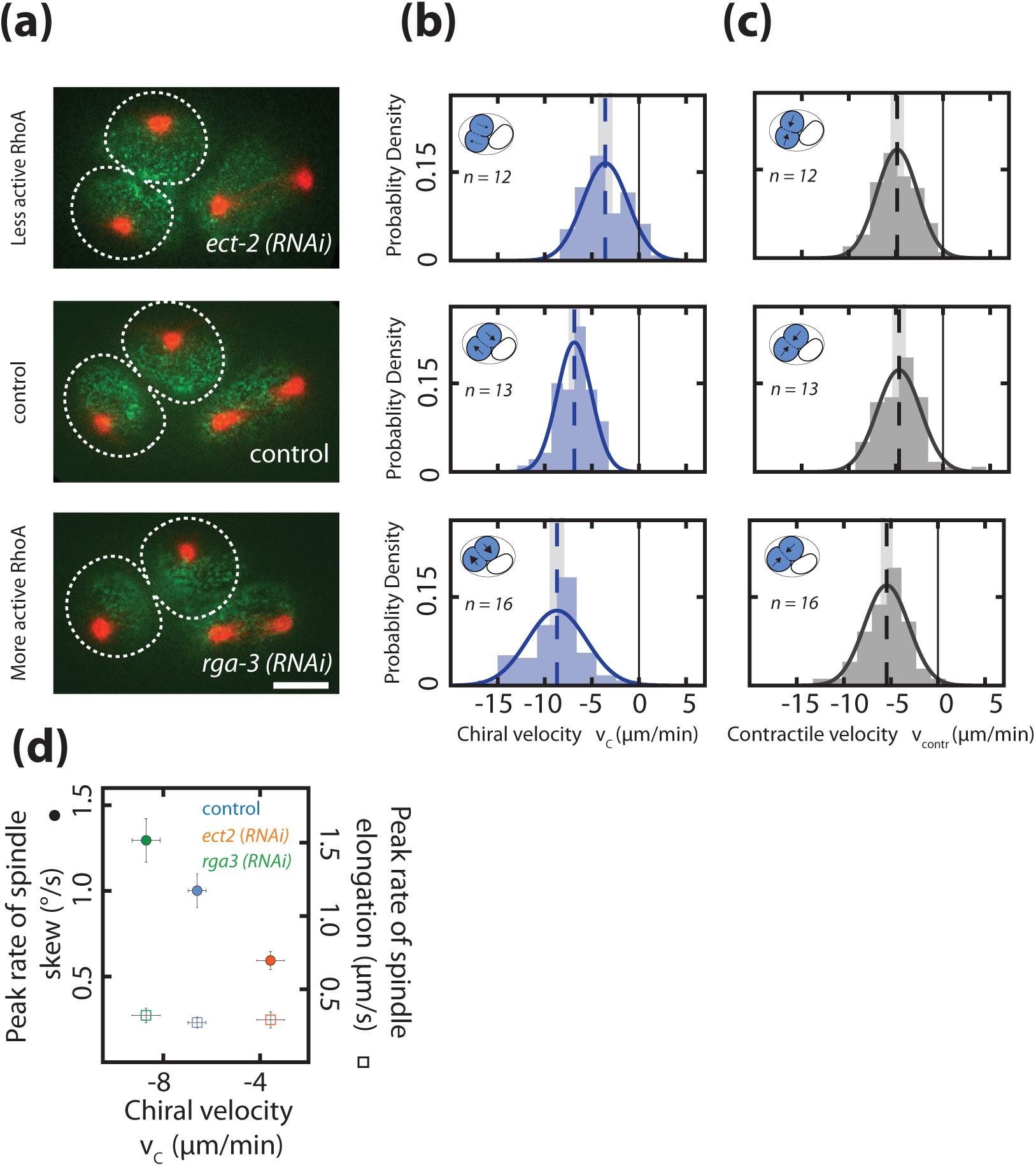
Chiral counter-rotating flow velocity determines the rate of spindle skew in the AB cell. **(a)**, Fluorescence images of two-cell embryos during cytokinesis upon *ect-2 (RNAi)* and *rga-3 (RNAi)*. Spindle poles are marked by TBB-2::mCherry (red), the cortex is marked by NMY-2::GFP (green). Dotted white lines indicate boundaries of the dividing AB cells. Scale bar, 10 *µ*m. **(b)**,**(c)**, Histograms of (**b**) the instantaneous chiral counter-rotating flow velocity v_c_ (see methods for definition) and **(c)** the instantaneous contractile flow velocity v_contr_, representing the flow velocity along the long axis of the dividing cell (see methods for definition). Dotted vertical colored lines represent the mean v_c_ or mean v_contr_, grey boxes represent the error of the mean. Thin black solid lines indicate zero velocities. Inset, blue shading indicates the AB cell analyzed, arrows indicate rotatory cortical flow. **(d)**, Peak rate of spindle skew (filled circles) and peak rate of spindle elongation (open squares) with the RNAi feeding control (control, *n = 10*; *ect-2 (RNAi), n=12, rga-3 (RNAi), n = 9*). Errors indicate the error of the mean at 95% confidence.

If chiral counter-rotating flows indeed drive spindle skews, we predict that a complete absence of chiral flows would lead to a complete absence of spindle skews. We test this prediction in the AB cell, by inactivating cortical flows entirely through a fast acting temperature sensitive *nmy-2(ts)* mutant [49]. Worms were dissected to obtain one-cell embryos and mounted at the permissive temperature at 15°C, to allow them to develop normally. Approximately 60 s after the completion of the first (P_0_) cell division, the temperature was rapidly shifted to the restrictive temperature of 25°C using a CherryTemp system. Imaging was started as soon as the AB cell entered anaphase. In all *nmy-2(ts)* mutant embryos (12 out of 12) an ingressing cytokinetic ring was absent and cytokinesis failed after temperature shifting, indicating that NMY-2 was effectively inactivated [50] (Supplement Video 14). Interestingly, while the dynamics of spindle elongation, including the peak rate of spindle elongation and the final spindle length, are essentially unchanged as compared to control embryos at restrictive temperature, the average peak rate of spindle skew was significantly reduced as compared to control embryos (0.33 ± 0.09°/min as compared 1.43 ± 0.13°/min in control embryos; Fig. 4(a)-(b); Supplement Video 13, 14). This demonstrates that a functional actomyosin cortex with chiral cortical flows is required for spindle skews of the AB cell. In light of our experiments above, we now conclude that spindle skews in the AB cell are independent of spindle elongation, but are instead driven by chiral counter-rotating flows. Finally, we performed temperature shift experiments for the first 11 divisions in the nematode, and found that all cells belonging to the AB cell lineage show a significantly reduced skew in the *nmy-2(ts)* embryos at restrictive temperatures as compared to the control (Fig. 4(c)). We conclude that a functional actomyosin cortex with chiral cortical flows is required for spindle skews of early AB lineage cell divisions during worm development. Furthermore, spindle skews and cell repositioning of early AB lineage cells are independent of spindle elongation, but are instead driven by chiral counter-rotating flows.

**Figure 4:**
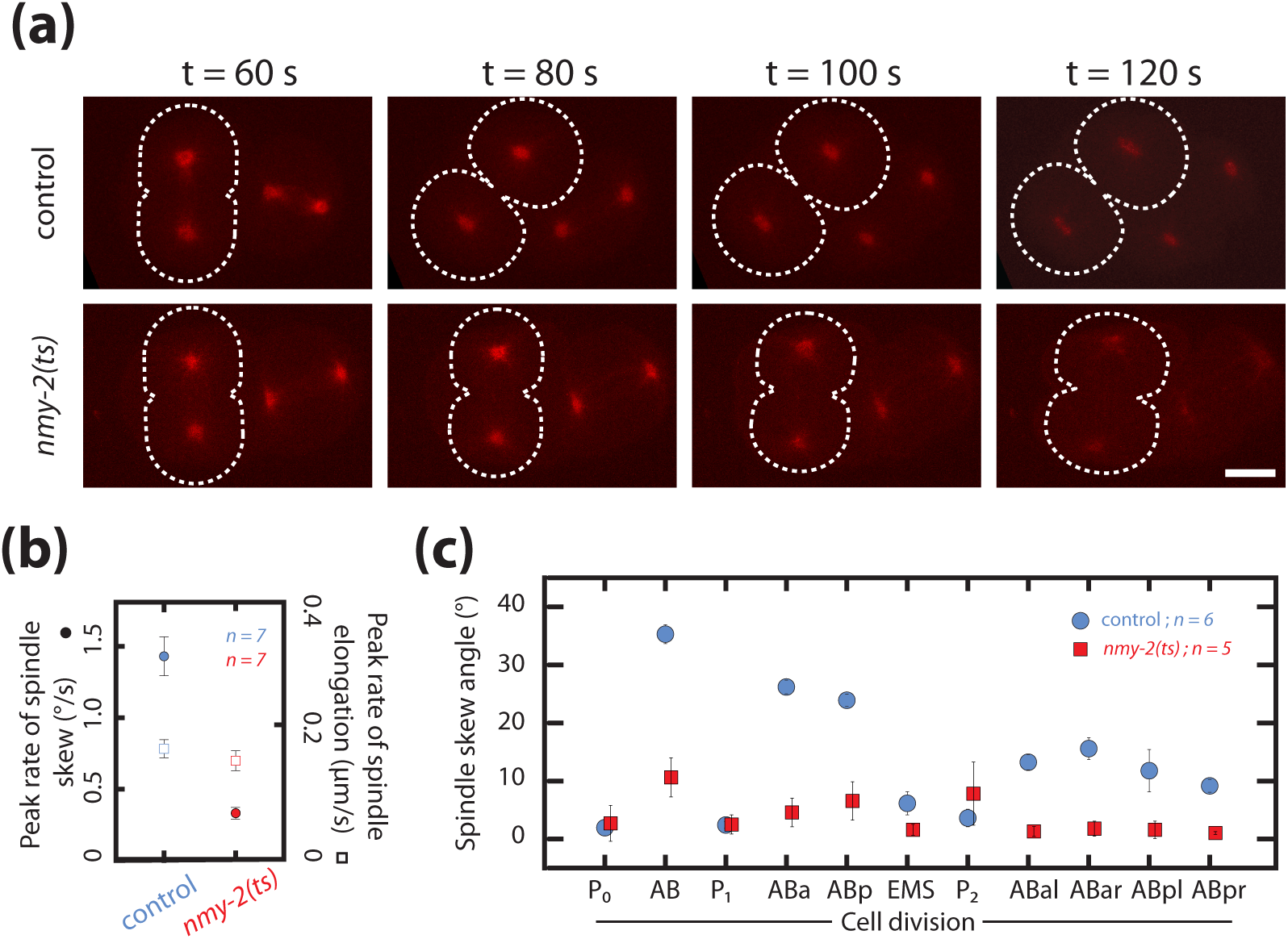
Spindle skews in early development are driven by chiral actomyosin flows. **(a)**, Fluorescence images of control and myosin temperature-sensitive (ts) mutants (*nmy-2(ts)*, permissive at 15°C, restrictive at 25°C) two-cell embryos with TBB-2::mCherry labelled for visualizing spindle poles (red). All embryos were shifted from 15°C to 25°C at the end of the first division. Dotted white lines indicate boundaries of the dividing AB cells. Time t = 60s indicates the onset of cytokinesis. Scale bar, 10 *µ*m. **(b)**, Corresponding peak rate of spindle skew (circles) and spindle elongation (squares) for the AB cell in control and *nmy-2(ts)* embryos. The full time series is given in Supplementary Fig. 4(d) and 4(e). **(c)**, Spindle skew angle in control embryos (blue circles) and *nmy-2(ts)* embryos (red squares) during the first 11 cell divisions at temperature of 25°C. Errors indicate the error of the mean at 95% confidence.

## DISCUSSION

Here, we show that lineage-specific chiral flows of the actomyosin cortex drive lineage-specific cell rearrangements in early *C. elegans* development. Recently, actomyosin flows were demonstrated to be required for determining the initial orientation of the mitotic spindle [17]. We here show that actomyosin cortex drives an active counter-rotation of the two halves of dividing early AB lineage cells, leading to cellular repositioning of these cells during cell division. We suggest that chiral counter-rotating flows represent a general mechanism by which dividing cells can reorient themselves in order to allow their daughters to achieve their appropriate positions.

How does the cell lineage determine the presence or absence of chiral cortical flows? We know that AB lineage cells divide symmetrically, while the early P/EMS lineage cell divisions are asymmetric [51]. Accordingly, we here show that cell divisions in the AB lineage are symmetric in terms of cytokinetic ring positioning and myosin distribution and display chiral counter-rotating flows, while divisions in the P/EMS lineage are asymmetric in these measures and chiral counter-rotating flows are absent (Fig. 1(b)-(e)). Cell lineage determinants like PAR proteins, instrumental for cell polarity, may therefore control chiral flows by modulating the distribution of molecular motors along the cell division axis [52]. In addition, many cortical constituents polarise along the anteroposterior axis prior to the first division, resulting in the AB cell to inherit higher levels of actin, myosin and its regulators [53–55] (Fig. 1(c)). The molecular composition of the cortex may therefore be different in the AB and P/EMS lineage, which is consistent with a recent study showing that both lineages are differentially dependent on the actin nucleator CYK-1/mDia for successful cytokinesis [56]. Together, the combined action of cortex composition and the intracellular distribution of molecular motors may also determine the type of chiral flow behavior (Supplement Fig. 2).

By which mechanism are active torques generated? We showed previously that chiral active fluid theory can quantitatively recapitulate the chiral counter-rotating flows as observed during *C. elegans* zygote polarization. In this coarse-grained hydrodynamic description, the cortex is treated as a two-dimensional gel that has liquid-like properties and generates active torques. Assuming that the local torque density is proportional to the measured NMY-2 levels, we here show that this theory can quantitatively recapitulate the observed chiral flow behaviour in the dividing AB blastomeres (Supplement Fig. 3(a)-(b)). However, given that many actin-binding proteins (e.g. actomyosin regulators, nucleators and cross-linkers) localise in a pattern similar to myosin, and many of these affect the chirality of actomyosin flows in the polarising one-cell embryo, we cannot exclude that other force generators provide the active torque needed for chiral counter-rotating flows. Interestingly, like myosins [41], diaphanous-like formins are known to generate torque at the molecular level [57, 58] and are required for numerous cellular [59] and organismal chiral processes [60–62]. Future work will be needed to first identify the molecular torque generator and to subsequently unravel the molecular mechanism underlying active torque generation. What is needed for driving cellular rearrangement by chiral counter-rotating flows, however, is an asymmetry in friction between the dividing cell and its environment (Fig. 2(f)). We speculate that the dividing cell simultaneously contacts different surfaces (e.g. neighboring cells and the eggshell), which gives rise to an inhomogeneous friction coefficient [17], thus allowing the cell to rotate and reposition.

To conclude, our work shows that chiral rotatory movements of the actomyosin cortex are more prevalent in development than one might have suspected. We report on an intricate pattern of chiral flows during early *C. elegans* development, leading to dedicated cell skew and cell reorientation pattern. This is interesting from a physical perspective, since together the cells of the early embryo represent a new class of active chiral material that requires an explicit treatment of angular momentum conservation [33, 44]. This is interesting from a biological perspective, since cell skews driven by actomyosin torque generation represent a novel class of lineage-specific morphogenetic rearrangements. Together, our work provides new insights into chiral active matter and the mechanisms by which chiral processes contribute to animal development.

## MATERIALS AND METHODS

### Worm strains and handling

*C. elegans* worms were cultured on NGM agar plates seeded with OP50 as previously described [63]. The following *C. elegans* strains were used in this study:

LP133 : *nmy-2(cp8[nmy-2::GFP + unc-119(+)]) I; unc-119(ed3) III* for imaging cortical flows [64].
SWG063 : *nmy-2(cp8[nmy-2::GFP + LoxP])I; weIs21[Ppie-1::mCherry::beta-tubulin::pie-1 3’UTR] IV; unc-119(ed3)III* (cross between (LP133 and JA1559 (Gift from J. Ahringer lab)) for measuring spindle skews in the AB cell.
SWG204 : *nmy-2(ne3409ts) I; unc-119(ed3)III; weIs21[pJA138(Ppie-1::mCherry::beta-tubulin::pie-1 3’UTR)] IV* (cross between WM179 and JA1559) for temperature sensitive experiments.
TH155 : *unc-119(ed3)III; weIs21[pJA138(Ppie-1::mCherry::beta-tubulin::pie-1 3’UTR)] IV; PH::GFP* (cross between JA1559 and OD70 [65]) for measuring spindle skews with the membrane marker and to measure relative cytokinetic ring position along the cell division axis.

### RNA interference

All RNAi experiments in this study were performed at 20°C [66]. For performing weak perturbation RNAi experiments, L4 staged worms were first incubated overnight on OP50 plates. Young adults were then transferred to respective RNAi feeding plates (NGM agar containing 1 mM isopropyl-*β*-D-thiogalactoside and 50 *µ*g ml-1 ampicillin seeded with RNAi bacteria). Feeding times for RNAi experiments were 4-5 hrs for *ect-2 (RNAi)* and 6-8 hrs for *rga-3 (RNAi)*. For *par-2, chin-1, lgl-1 (RNAi)* and *par-6 (RNAi)* conditions, young L4 were transferred to feeding plates 24 hrs before imaging. Embryos used as controls for all RNAi experiments were grown on plates seeded with bacteria containing a L4440 empty vector, and exposed for the same number of hrs as the corresponding experimental RNAi perturbed worms.

The indicated hrs of RNAi treatment reflects the time that worm spent on the feeding plate. All dissections were done in M9 buffer to obtain early embryos. Embryos were mounted on 2% agarose pads for image acquisition for the all cell divisions except 4-6 cell stage [67]. Four cell embryos were mounted using low melt agarose [31, 33]. The *rga-3* feeding clone was obtained from the Ahringer lab (Gurdon institute, Cambridge, United Kingdom) and the *ect-2* feeding clone from the Hyman lab (MPI-CBG, Dresden, Germany). RNAi feeding clones for *par-2, par-6, chin-1 and lgl-1* were obtained from Source Bioscience (Nottingham, United Kingdom).

### Image Acquisition

All the imagining done in this study was performed at room temperature (22-23°C) with a spinning disk confocal microscope. Two different spinning disk microscope systems were used for the study.

### System 1

Axio Observer Z1 - ZEISS spinning disk confocal microscope; Apochromat 63X/1.2 NA objective lens; Yokogawa CSU-X1 scan head; Andor iXon EMCCD camera (512 by 512 pixels), Andor iQ software.

### System 2

Axio Observer Z1 - ZEISS spinning disk confocal microscope; Apochromat 63X/1.2 NA objective lens; Yokogawa CSU-X1 scan head; Hamamatsu ORCA-flash 4.0 camera (2048 by 2048 pixels); micromanager software [68].

Confocal videos of cortical NMY-2::GFP for the first three cell divisions for control and RNAi conditions were acquired using system 1. A stack was acquired using 488 nm laser and an exposure of 150 ms was acquired at an interval of 3s between image acquisition consisting of three z-planes (0.5 *µ*m apart). The maximum intensity projection of the stack at each time point was then used for further analysis. 21s of time frames were analyzed for each embryo starting from the time frame when the cytokinetic ring has ingressed ∼10 %.Time intervals between consecutive frames were increased at later cell stage to avoid photo-toxicity and bleaching. From the 4-6 cell stage onwards, imaging was performed using system 2. A time period of 20s was analyzed for measuring chiral counter-rotation flow velocity. Image stacks with a spacing of 0.5 *µ*m for 20 z-slices from the bottom surface towards the inside of the embryo were taken with 488 nm laser with exposure of 150 ms at an interval of 5 s during the time of cytokinetic ring ingression and projected with maximum intensity projection.

Imaging of the spindle poles (TBB-2::mCherry) was done using SWG63 (Fig. 2(e), 3(a), 3(d), 4(a)-(c)). TH155 strain was used to image TBB-2::mCherry together with PH::GFP to determine cytokinetic ring position and spindle skew angle in Fig. 2(b) for 11 cell stages. SWG204 strain was used for temperature sensitive experiments (Fig. 4(a)-(c)). Spindle pole imaging for AB cell (TBB-2::mCherry) was performed by acquiring nine-z-planes (1 *µ*m apart) with a 594 nm laser and exposure of 150 ms on the second system at an interval of 3s. For long term imaging using TH155 strain (Fig. 2(b)), TBB-2::mCherry together with PH::GFP was imaged in 24 z-slices (1 *µ*m apart) with 594 nm laser and exposure of 150 ms at an interval of 30 s (Supplement Video 9).

### Image Analysis

Cortical flow velocity fields in the xy-plane were obtained using a MATLAB code based on the freely available Particle Image Velocimetry (PIVlab) MATLAB algorithm [69]. Throughout the study, we used the 3-step multi-pass with linear window deformation, a step size of 8 pixels and a final interrogation area of 16 pixels.

To determine the average flow fields that were fitted by the chiral thin film theory (Supplement Fig. 3(a)), we first tiled the flow field determined by PIV into 9 sections along the long-axis of cells. Within each section, we averaged the flow field over the entire duration during which flows occur. Since we were only interested in flows that are generated by the cytokinetic ring, we excluded embryos in which we could not differentiate whole body rotation from cytokinesis [43]. From all the imaged cell divisions, a small subset of measurements in which the optical focus on the cortical plane was lost during imaging, was discarded.

From the measured cortical flows, we extracted properties that are later used to quantify their chiral counter-rotating nature in the following way. First, we identified the cleavage plane or the cytokinetic ring by eye. We then defined a region of interest (ROI) on each side of the cytokinetic ring (Fig. 1(a)). In order to ensure that PIV measurements were not affected by the saturating fluorescence signal from the cytokinetic ring, we placed the inner boundary of each ROI at a distance of 1 *µ*m from the ring. The position of the outer boundary of each ROI (along the long axis) was scaled with the size of the analyzed cells (10 *µ*m, 8.5 *µ*m, 6 *µ*m, 6.5 *µ*m, 6.5 *µ*m, 6.5 *µ*m for P_0_, AB, P_1_, ABa, ABp and EMS, respectively, measured from the position of the cytokinetic ring). For P_2_, ABal and ABpr cells the distance of the outer ROI boundary to the cytokinetic ring varied around 4 *µ*m, depending on the cell surface area visible in the imaging focal plane. For every cell, the width of ROIs was set equal to the length of the cytokinetic ring visible in the focal plane. In each ROI, we averaged the observed flow field over a period of 21 s. Consistent with earlier findings, we observed a whole cell rotation prior to P_0_ cytokinesis [43, 70]. We discarded a small subset of embryos (4 out of 17), where this whole cell rotation coincided with the time frames in which we performed the flow analysis.

In order to quantify chiral flows, we defined a chiral counter-rotating flow velocity v_c_ together with a handedness as follows. First, we introduce a unit vector that is orthogonal to the contractile ring and points towards the cell pole that is in the direction of the anteroposterior-axis of the overall embryo (<u>e_x_</u>, see Supplement Fig. 1(a)). A second unit vector <u>e_y_</u> is used, which is parallel to the cytokinetic ring and orthogonal to <u>e_x_</u>. We denote the average velocities determined in the two halves of the cell (measured within the ROIs described above) as <u>v_1_</u> and <u>v_2_</u>. With this, we define the chiral counter-rotating flow velocity v_c_ by

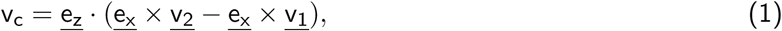

where <u>e_z_</u> = <u>e_x_</u> × <u>e_y_</u> is the third base vector of a right-handed orthonormal coordinate system. With this definition, |v_c_| represents the speed of chiral counter-rotating flows, while the sign of v_c_ determines the flow’s handedness: If cortical flows appear clockwise when viewed viewed from the cytokinetic plane towards each pole, one has v_c_ > 0 and the counter-rotating flows are left-handed. If cortical flows appear counter-clockwise, one has v_c_ < 0 and counter-rotating flows are right-handed.

To characterize net-rotating flows in a similar fashion, we use the net-rotating flow velocity v_r_ defined by (see Supplement Fig. 2(a))

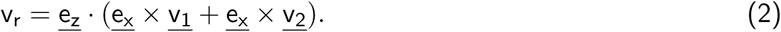

If average flows in both cell halves are equal and in the same direction, i.e. <u>v_1_</u> = <u>v_2_</u>, there are no counter-rotating flows (v_c_ = 0) and the corresponding net-rotating flow is quantified by v_r_.

Finally, in order to determine a flow measure that captures cortical flows along the cell’s long axis and into the contractile ring, we define the contractile flow velocity (v_contr_) depicted in Fig. 3(b) by

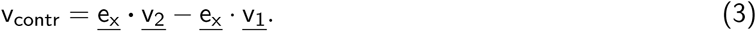

For contractile cortical flows into the cytokinetic ring, we have <u>e_x_</u> · <u>v</u>_1_ > 0 and <u>e_x_</u> · <u>v</u>_2_ < 0, and therefore vcontr < 0.

### Spindle skew and elongation analysis

In order to measure spindle skew angles, z-stacks (nine-planes; 1*µ*m apart at 3s time interval) of the spindle poles were first projected on the plane perpendicular to the cytokinetic ring. For P_0_, AB and P_1_ the imaging plane was already perpendicular to the plane of cytokinesis while for the remaining cell divisions the embryos were rotated accordingly using the clear volume plugin [71] in FIJI. Subsequently, the spindle skew angle was defined as the angle between the line that joins the spindle poles at the onset of spindle elongation (anaphase-B) (initial spindle angle) and at completion of cytokinesis (final spindle angle) (Fig. 2(a)). Spindle elongation length was defined as the distance between the two spindle poles during the same time-frames. To measure the dynamics of spindle elongation and skew angle in the AB cell (Supplement Fig. 4(b)-4(e)) the spindle poles were automatically tracked and the spindle skew angle was plotted using custom-written MATLAB code. In order to compare the dynamics of spindle elongation and spindle skew we first synchronized all the AB cells in each condition at the onset of anaphase-B and, subsequently, averaged over multiple embryos. The difference beween the spindle angle at each timepoint and the spindle angle at anaphase-B onset timepoint was measured and plotted over time. The maximum value of the slope for spindle elongation and spindle skew angle curves for each embryo between time window of 60-120s (see Supplement Fig. 4(b)-(e)) was identified as peak rate of spindle elongation and peak rate of spindle skew. Average peak rates were calculated by averaging the maximum values over multiple embryos for each condition.

### Determining myosin ratio, cytokinetic ring position and fitting myosin profiles

In order to obtain cortical myosin distributions (Fig. 1(c)) we normalized the axis of cell division and calculated the mean myosin intensity in a 20 pixel-wide stripe along this axis for the same timepoints when v_c_ was calculated. The myosin ratio (Fig. 1(e)) was determined by dividing the mean intensity at the anterior (0-20% of the normalized cell division axis) by the mean intensity at the posterior (80-100% of the normalized cell division axis) for P_0_, AB, P_1_ and EMS. For ABa and ABp the myosin intensity was calculated similarly but along the L/R axis. To determine the position of the cytokinetic ring, we performed midplane imaging using a strain expressing a membrane marker (PH::GFP) and a tubulin marker to label spindle poles (TBB-2::mCherry). The cell division axis was defined as the line through the two opposing spindle poles and was normalized between cell boundaries. The relative cytokinetic ring position was defined as the position where the ingressing ring intersects the normalized cell division axis.

For visualization purposes (Fig. 1(c)), we fitted the myosin distributions using the fitting function

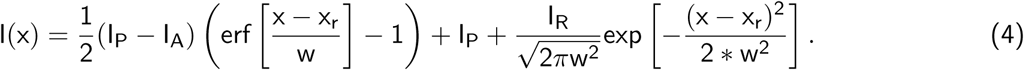

The errorfunction erf and the Gaussian represent contributions from the anteroposterior myosin asymmetry and from the contractile ring, respectively. The fitting parameters I_P_, I_A_, I_R_, w and x_r_, respectively, describe myosin intensities at the posterior pole, the anterior pole and in the cytokinetic ring, as well as the width and position of the ring.

### Temperature sensitive experiments

We used the cherry temp temperature control stage from Cherry Biotech for all temperature sensitive experiments in the study. L4 worms carrying *tbb-2::mCherry* (SWG63, control) and *nmy-2(ts); tbb-2::mCherry* (SWG204) were grown at 15°C (permissive temperature for *nmy-2(ts)* mutants) overnight. During image acquisition temperature was maintained using the cherry temp temperature control stage from Cherry Biotech. For imaging the dynamics of spindle elongation and skew angle in the AB cell for *nmy-2(ts)* and control embryos, 60 s after the successful completion of the first cell division (P_0_ cell division), temperature was raised to 25°C. Embryos were imaged approximately 5 min after the temperature shift to minimize photo toxicity. Similarly, for later cell stages, the temperature shift was carried out right after completion of cytokinesis of the mother cell. A small subset of embryos in which spindle poles of daughter cells could not be visualized were discarded.

## OTHER DETAILS

### Author Contributions

L.P., T.C.M., A.M. and S.W.G. conceptualized the work, analyzed the data and wrote the manuscript together. L.P. performed experiments with help from T.C.M. A.M. developed the theory with help from all authors.

## Acknowledgements

We thank Masatoshi Nishikawa for critical inputs on the theory, A. Honigmann and P. Gönczy for critical reading of the manuscript. We also would like to thank T. Hyman and B. Bowerman for sharing *C. elegans* strains and all the members of Grill lab for critical input and discussions. Some strains were provided by the CGC, which is funded by NIH Office of Research Infrastructure Programs (P40 OD010440). Furthermore, we thank the light microscopy facility of CMCB of TU Dresden, and the light microscopy facility of MPI-CBG for their support. We would like to acknowledge a long standing collaboration with Frank Jülicher on active chiral matter, and thank him for many discussions that were essential for the work here. L.P. acknowledges funding from the European Union’s Horizon 2020 research and innovation program under the Marie Sklodowska-Curie grant agreement No 641639. T.C.M was supported by the European Molecular Biology Organization (EMBO) long-term fellowship ALTF 1033-2015, and by the Dutch Research Council (NWO) Rubicon fellowship 825.15.010. A.M. thanks the ELBE Phd fellowship from the Center for Systems Biology, Dresden. S.W.G was supported by the DFG (SPP 1782, GSC 97, GR 3271/2, GR 3271/3, GR 3271/4), the European Research Council (grants 281903 and 742712).

## Competing Interests

The authors declare that they have no competing financial interests.

## SUPPLEMENTARY DATA

### Thin film active chiral fluid theory

Cortical flows in *C. elegans* embryos have been successfully described using the thin film theory of active chiral fluids [33, 40]. We use the fact that cells are approximately axisymmetric, and we assume that the active forces during cell division vary mostly along their symmetry axis, in the following referred to as *long axis*. In this case, the hydrodynamic equations of a chiral thin film can be formulated as an effectively one-dimensional system of equations, capturing flows parallel (v_x_) and orthogonal (v_y_) to the long axis:

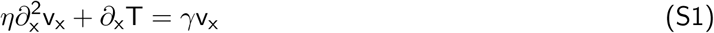

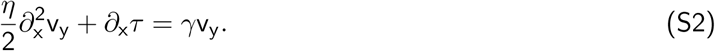

Here, *η* is the viscosity of the cortex and *γ* is the friction with surrounding material. Note that the assumption of a homogeneous friction *γ* is a strong simplification, where we neglect details of the potentially inhomogeneous mechanical interactions between cells, as well as with the extra-embryonic fluid and the egg shell. Gradients of active tension T and active torques *τ* lead to contractile and chiral flows, respectively. It has been demonstrated previously that active tension and active torques both dependent on the local myosin concentration in the cortex [33, 35, 40]. Using the fluorescent myosin intensity I(x) as a proxy for the myosin concentration, we can write in the simplest case T = T_0_I(x) and T = *τ*_0_I(x).

Denoting the length of the measurement domain by L (approximately the length of the cell’s long axis), we can identify three independent parameters in the model Eqs. (S1) and (S2): The characteristic contractile and chiral velocities v_T_ = LT_0_*/η* and v_*τ*_ = L*τ*_0_*/η*, respectively, as well as the hydrodynamic length 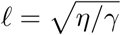.

### Fitting cortical flows

For the fitting of cortical flows for given myosin profiles we closely follow our previous work [33]. Briefly, we use experimentally determined myosin profiles I(x) in Eqs. (S1) and (S2) to calculate flows v_x_ and v_y_. We then determine the parameters v_T_, v_*τ*_ and *𝓁* for which the predicted flows best match the experimental data. To extract unique solutions from Eqs. (S1) and (S2), we use the experimentally measured flow velocities at the boundaries of the measurement domain as boundary conditions. Note that due to imaging limitations for smaller cells (after the 8-cell stage) as development progresses, it was not possible to extract spatially resolved velocity line profiles and fits for the cortical flows in ABal cell and ABpr cell. The final best fit parameters for AB, ABa and ABp cells are shown in Tab. S1. The corresponding contractile flow profiles v_x_ and chiral flow profiles v_y_ are shown in Supplement Fig. 3(a) and 3(b).

**TABLE S1:** Best fit parameters. The average domain lengths (approximately the size of the cell’s long axis) on which the myosin profiles were given were L_AB_ = 27.4 *µ*m, L_ABa_ = 19.5 *µ*m and L_ABp_ = 17.9 *µ*m.

| Cell type | $\ell$ [$\mu\text{m}$] | $v_T$ [$\mu\text{m}/\text{min}$] | $v_\tau$ [$\mu\text{m}/\text{min}$] |
| --- | --- | --- | --- |
| AB | 18.7 | 0.15 | 0.25 |
| ABa | 10.4 | 0.19 | 0.15 |
| ABp | 8.2 | 0.4 | 0.4 |

We also considered asymmetrically dividing cells (P_0_, P_1_, EMS) in which, instead of counter-rotating flows (|v_c_| > 0, v_r_ ≈ 0), mainly net-rotating flows occur (v_c_ ≈ 0, |v_r_| > 0) (Supplement Fig. 2). Following the same fitting procedure as described in the previous section, we noticed that the chiral thin film theory Eqs. (S1) and (S2) generally could not account quantitatively for the observed cortical flow profiles. However, using the theory it is still possible to rationalize qualitative properties of the observed cortical flows, as we discuss in the following.

### Qualitative properties of myosin distributions and chiral flows

While the chiral thin film theory does not recapitulate all of the experimentally observed flows quantitatively, the theory allows linking qualitative predictions to key properties of observed myosin distributions and chiral flows. In particular, we noticed that the myosin profiles in asymmetrically diving cells consistently featured a rather asymmetrically positioned contractile ring (Fig. 1(d)). Furthermore, the myosin profiles of asymmetrically dividing cells are asymmetric with respect to the cytokinetic ring and exhibit plateaus towards the anterior cell poles (Fig. 1(c) and Fig. 1(e)). An overview of these qualitative properties for the P/EMS lineage and the AB-lineage is given in Table S2.

**TABLE S2:**
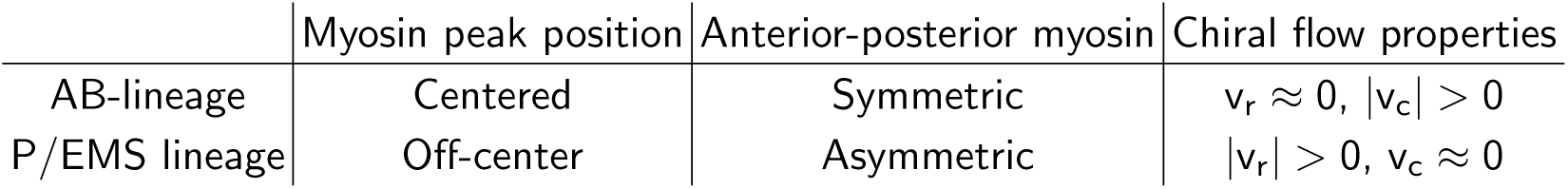
Qualitative properties of myosin distributions and chiral flows consistently observed in P/EMS lineage (P_0_, P_1_, EMS, P_2_) and AB-lineage (AB, ABa, ABp, ABal, ABar) cells.

To establish how a contractile ring and an overall myosin asymmetry generally affect flows predicted by the chiral thin film theory, we generalize the chiral thin film equations [33, 40] to curved surfaces and solve them on an ellipsoid using corresponding synthetic torque profiles (Supplement Fig. 3(c)). We find that a symmetrically placed ring pattern yields perfectly counter-rotating flows (v_r_ = 0, |v_c_| > 0, Supplement Fig. 3(c), left), while an anterior-posterior myosin asymmetry yields net-rotating flows (|v_r_| > 0, v_c_ = 0, Supplement Fig. 3(c), right). To study the combined effect of a varying position of the contractile ring and a myosin asymmetry, we develop in the following section a simple minimalistic model of the system.

### Chiral flow minimal model

In the following, we develop a minimal description of the occurrence of counter- and net-rotating flows based on key characteristics of the myosin profiles discussed in the previous section. We consider a simplified myosin profile in the form

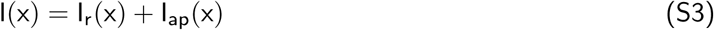

with

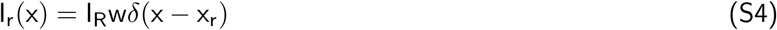

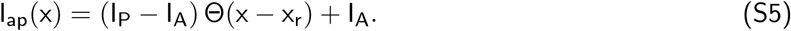

Here, *δ*(x) is the delta function, Θ(x) the step function and x_r_ is the position of an idealized cytokinetic ring. The intensity profile I_r_(x) captures the presence of a myosin peak (the later contractile ring) of width w and average intensity I_R_ at x_r_. The profile I_ap_(x) represents the large-scale asymmetry in myosin between anterior pole (I_A_) and posterior pole (I_P_). We solve Eq. (S2) on a domain of length L and for boundary conditions v_y_(0) = v_y_(L) = 0 using the simplified myosin intensities given in Eqs. (S4) and (S5), which yields

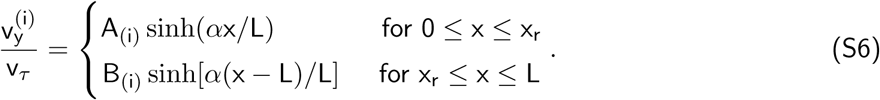

Here, 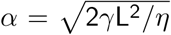 is an inverse dimensionless hydrodynamic length and v_*τ*_ = L*τ*_0_*/η* the characteristic velocity associated with active torques. The coefficients in Eq. (S6) are given by

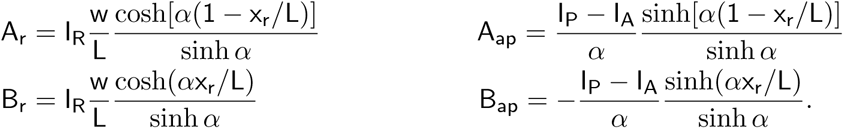

With these coefficients, Eq. (S6) describes flows 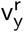 resulting from the contractile ring given in Eq. (S4) and flows 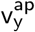 resulting from the asymmetric myosin profile given in Eq. (S5). Note that, as expected, flows due to the presence of the contractile ring vanish for I_R_ = 0 and flows due to the presence of a myosin asymmetry vanish if IA = I_P_. General flow profiles are given as superposition of the two contributions: 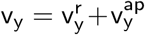.

Finally, to evaluate these solutions, we define a dimensionless counter-rotating velocity 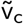 and the net-rotating velocity 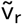 in analogy to the quantities introduced in the main text as

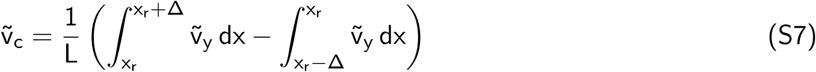

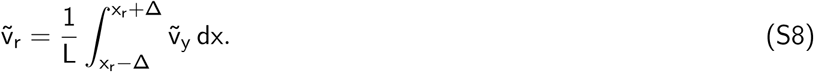

Here, 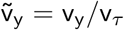 and we consider a window of 15 % (Δ/L = 0.15) of the total length towards either side of the idealized contractile ring over which the mean flow velocity is determined (Fig. 1(a), Supplement Fig. 1(a)). Using this minimal model, we can now investigate how the combined contributions of a large-scale myosin asymmetry (I_ap_) and a varying contractile ring position (I_r_) affect the counter-rotating velocity 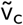 and net-rotating velocity 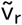. In particular, we fix for the discussion *α* = 1 and w/L = 0.1, as well as I_R_/I_P_ = 2. In this case, the counter-rotating flow velocity 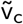 and the net-rotating flow velocity 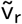 are only functions of the relative ring position x_r_/L and the anterior-posterior myosin ratio I_A_/I_P_ with properties shown in Supplement Fig. 3(d)-(g) and described in the following.

For an anterior-posterior symmetric myosin profile, counter-rotating flows 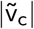 have a weak maximum in their amplitude if the contractile ring is at a centered position x_r_/L = 0.5 (Supplement Fig. 3(d)). Furthermore, a centered contractile ring has no effect on the counter-rotating flow measure 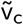 if a large-scale myosin asymmetry I_A_/I_P_ > 1 is introduced (Supplement Fig. 3(e)). However, for an off-centered ring x_r_/L > 0.5 the presence of a myosin asymmetry will contribute flows that reduce 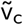 and can even lead to vanishing counter-rotating flows 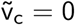 if I_A_/I_P_ is sufficiently large. While this matches the qualitative experimental observations listed in Tab. S2, the required asymmetry predicted by the theory is significantly larger than the experimentally measured one.

Also, the behavior of 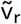 in our minimal model is compatible with the qualitative properties listed in Tab. S2. In particular, for a large-scale myosin asymmetry I_A_/I_P_ > 1, the net-rotation velocity 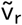 is negative for any ring position, while it can becomes positive for x_r_/L > 0.5 if I_A_ = I_P_ (Supplement Fig. 3(f)). Furthermore, net-rotating flows vanish for a symmetric myosin profile I_A_ = I_P_ and a centered contractile ring x_r_/L = 0.5 (Supplement Fig. 3(g)), which corresponds to the observations made in AB-lineage cells. Finally, the presence of large-scale myosin asymmetries I_A_/I_P_ ≠ 1 is generally expected to contribute to net rotating flows (Supplement Fig. 3(g)). This holds true for essentially arbitrary positions of the contractile ring and indicates that such asymmetries could play an important role for developing a better understanding of net-rotating flows in the P/EMS lineage cells in the future.

## SUPPLEMENTARY FIGURE LEGENDS

**Supplement Fig. 1:**
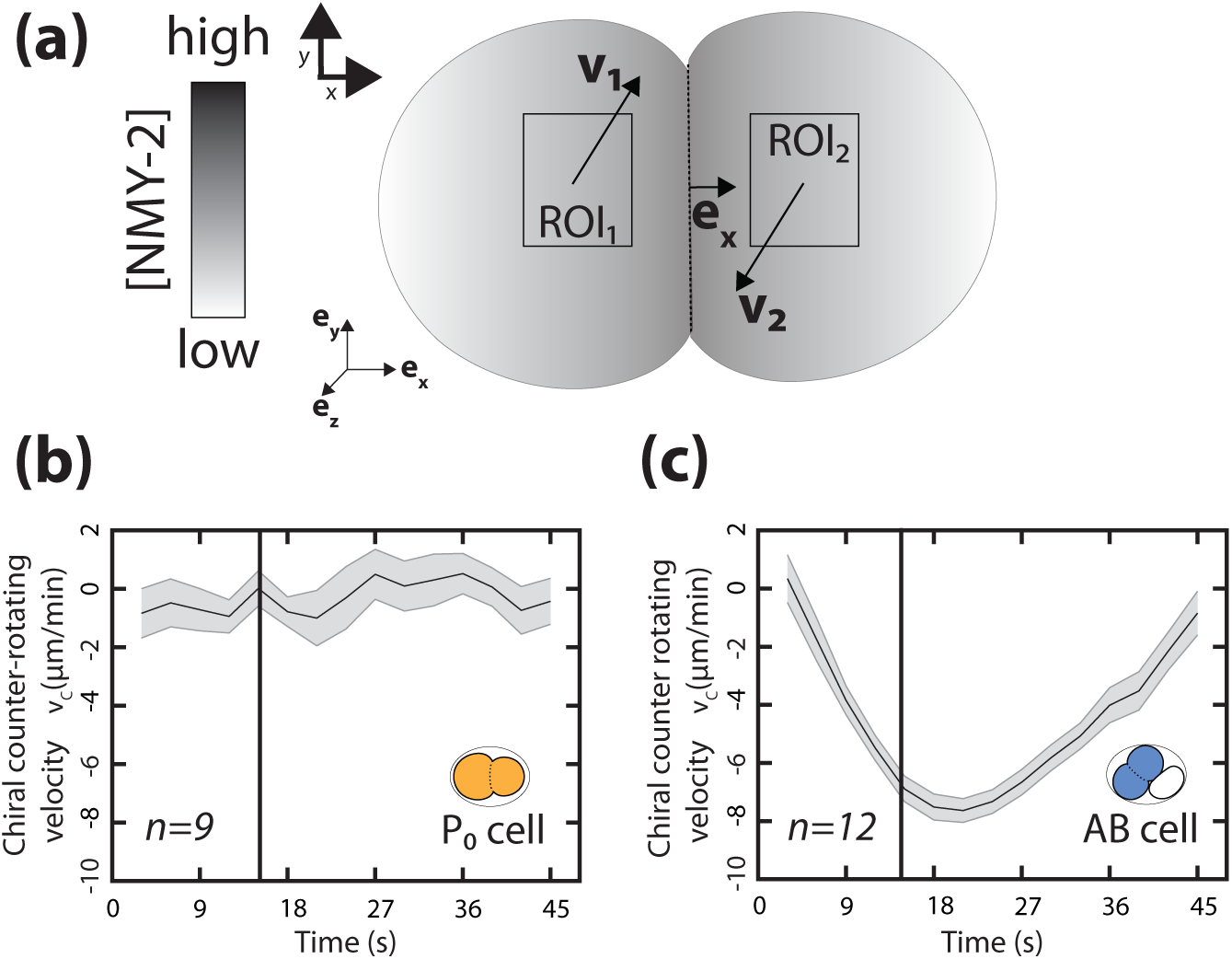
Time evolution of chiral counter-rotating velocity during cytokinesis. **(a)**, Schematic illustrating the measured chiral counter-rotating velocities in a dividing cell. v_c_ = <u>e_z_</u> · (<u>e_x_</u> × <u>v_2_</u> − <u>e_x_</u> × <u>v_1_</u>). Here <u>e_x_</u>, <u>e_y_</u> and <u>e_z_</u> are orthonormal vectors depicted in left corner. In this definition, the magnitude of v_c_ gives us the speed of the flows and the sign of v_c_ denotes the handedness of chiral counter-rotating flows. In this sketch, the counter-rotating pair has right-handedness as observed in the AB lineage (see methods for more information on definition of handedness and v_c_; upper left indicates the coordinate system).**(b), (c)**, Chiral counter-rotating flow speed (v_c_) measured over seven subsequent frames for different stages for **(b)** the P_0_ and **(c)** the AB cell from the beginning of cytokinesis until completion of cytokinesis. Vertical black line marks the 10% cytokinetic furrow ingression, used as a reference timepoint to measure v_c_. The curve represents the mean v_c_, grey shaded bar indicates error of the mean at 95% confidence interval. Inset, colored cell indicates the cell analyzed.

**Supplement Fig. 2:**
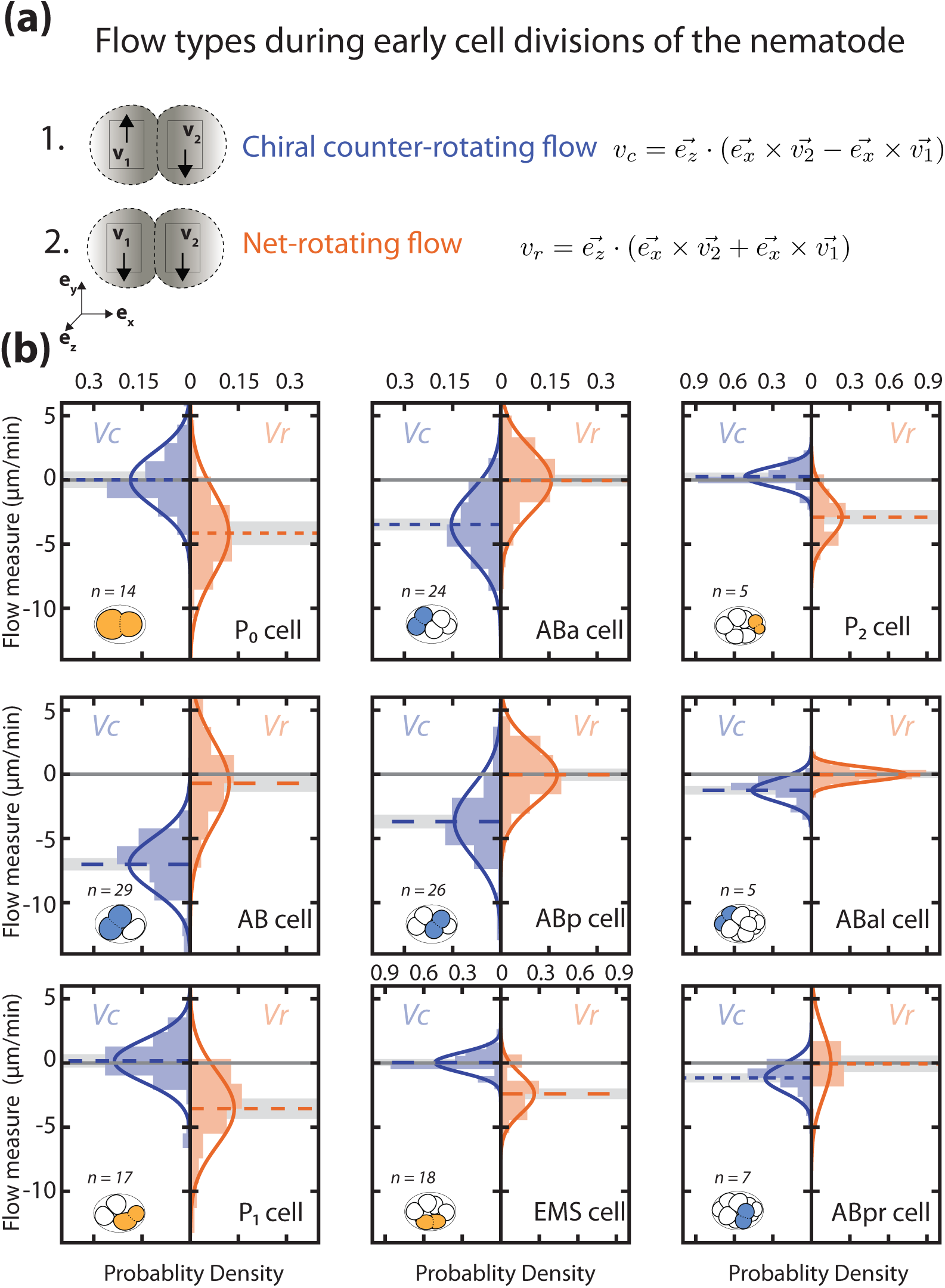
Early cell divisions display different types of rotatory flows. **(a)**, Schematics of chiral counter-rotating flows (upper part) and net-rotating flows (lower part) observed in the AB and P/EMS lineage respectively. Arrows mark flow direction. When the rotatory flows in the two halves of the dividing cell counter-rotate, v_c_ is maximum in magnitude and v_r_ is minimum in magnitude. This type of flow is seen in the case of cells of AB lineage. However when rotatory flows in the two halves of the cell rotate in the same direction, v_c_ is minimum and v_r_ is maximum. Such a flow is observed in the case of cells of the P/EMS lineage. **(b)**, Histograms of the instantaneous chiral counter-rotating flow velocity v_c_ (left; blue) and instantaneous net-rotating velocity v_r_ (right; orange) plotted for first nine cell divisions. v_c_ and v_r_ are calculated with denoted equations in **(a)** (see Supplement Fig. 1(a) and methods for more information). Solid lines indicate the best-fit gaussian probability density function. Dotted colored lines represent the mean v_c_ on the left and mean v_r_ on the right, grey shaded bar represent the error of the mean at 95% confidence interval. Thin black solid lines indicate a chiral flow velocity of zero. Inset, colored cells indicate the cell analyzed, AB lineage in blue and P/EMS lineage in orange.

**Supplement Fig. 3:**
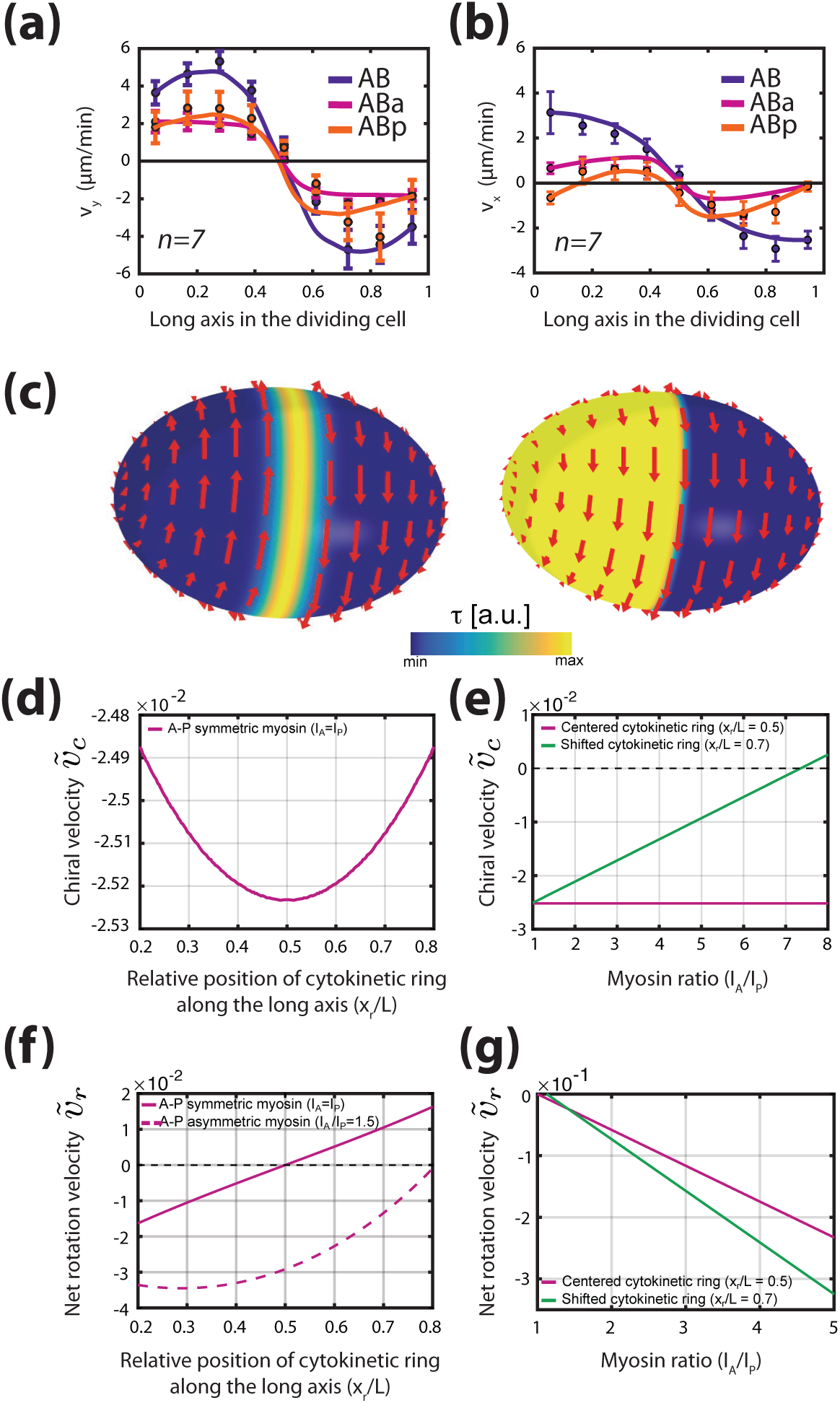
Active chiral fluid theory can recapitulate the contractile and rotatory flows observed in the cells of the AB-lineage. **(a), (b)**, Velocity profiles (black filled circles) along the long axis of the dividing cell and theoretical prediction (solid line) for (**(a)**) chiral counter-rotating flow velocity and (**(b)**) contractile on-axis flow velocity in the cells belonging to the AB lineage (blue, AB; magenta, ABa; orange, ABp) obtained from thin film active chiral fluid theory using NMY-2::GFP gradients measured during cytokinesis. For the AB cell, a representative NMY-2::GFP gradient is shown in Fig. 1(c). **c**, Active chiral thin film model Eq. (S2) solved on an ellipsoidal surface for two different profiles of active torques. Left: A centered ring of active torques (described by a Gaussian of width w/L_s_ = 0.1, where L_s_ is the meridional length on the surface from one pole to another. Active torques organized in this fashion lead to counter-rotating flows. Right: Pole-to-pole asymmetry of active torques described by an error function with step width w/L_s_ = 0.1 (see Eq. (4) methods). Such a large-scale asymmetry leads to net-rotating flows. Profile of active torque generator is shown in the blue-to-yellow shading. **(d)-(g)**, Properties of counter-rotating and net-rotating flows in a minimal model (Fixed parameters: *α* = 1, w/L = 0.1, I_R_/I_P_ = 2). **(d)**, Chiral counter-rotating velocity 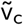 for a symmetric myosin distribution I_A_ = I_P_ and varying position of the contractile ring. 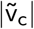 has a weak maximum for a centered contractile ring at x_r_/L = 0.5. **(e)**, Chiral counter-rotating velocity 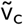 for varying asymmetry of myosin between anterior and posterior poles. While 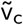 is independent of any myosin asymmetry for a centered contractile ring (magenta curve), counter-rotating flows may vanish for sufficiently large myosin asymmetries, if the contractile ring is positioned off-center (green curve) **f**, Net-rotating flows velocity 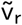 for varying position of the contractile ring. If I_A_ = I_P_, net-rotating flows 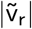 vanish for a centered contractile ring (solid line). In the presence of a sufficiently large anterior-posterior myosin asymmetry I_A_ > I_P_ one finds 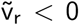 for any ring position (dashed curve). **(g)**, Net-rotating velocity 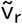 for varying asymmetry of myosin between anterior and posterior poles. Irrespective of the contractile ring position, myosin asymmetries (here shown for I_A_ > I_P_) generally contribute to net-rotating flows.

**Supplement Fig. 4:**
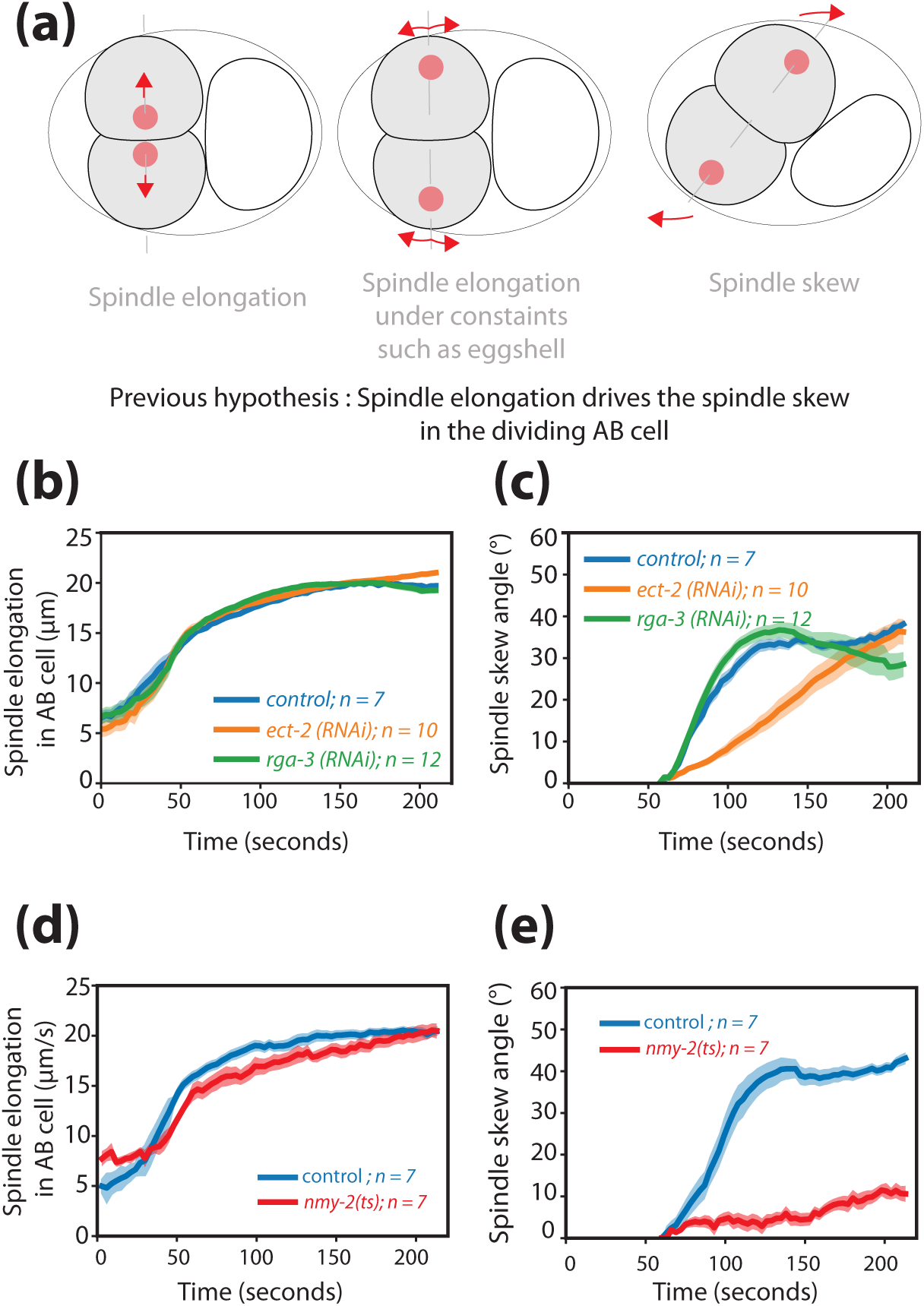
Spindle elongation and spindle skew dynamics during cytokinesis in the AB cell. **(a)**, Illustration of the previously hypothesized mechanism where spindle elongation drives spindle skew in the AB cell. The anterior positioned AB cell (cell on the left) divides before the posterior P_1_ cell (cell on the right). The spindle initially elongates along the dorsoventral axis (left and middle illustrations) and subsequently skews (right illustration) due to constraints imposed by the egg shell. Dashed line depicts the cell division axis, red dots depict spindle poles, arrows indicate spindle pole movements. **(b), (c)**, Dynamics of (**(b)**) spindle elongation length and (**(c)**) spindle skew angle in the AB cell during cytokinesis for RNAi feeding control and *ect-2(RNAi)* embryos and *rga-3(RNAi)* embryos. The dynamics of spindle length in these perturbations is not affected refuting the first hypothesis as depicted in (**a**) and supporting the alternative hypothesis proposed here, as illustrated in Fig. 2g (see main text). **(d), (e)**, Dynamics of (**(d)**) spindle elongation length and (**(e)**) spindle skew angle in the AB cell during cytokinesis for *tbb-2::mCherry* (control) and *nmy-2(ts); tbb-2::mCherry* embryos at 25°C. Solid line represents mean whereas the shaded region is error of the mean at 95% confidence.

## Supplementary Videos

**Video 1**: Movie of the actomyosin cortex during the first cell division.The cortex is marked by NMY-2::GFP. The P_0_ cell does not exhibit chiral counter-rotating flows during its division. Cortical flows in the two dividing halves of the P_0_ cell flow in the same y-direction. Scale bar 10*µ*m.

**Video 2**: Movie of the actomyosin cortex during the second cell division. The AB cell division exhibits chiral counter-rotating flows. The two dividing halves of the AB cell counter rotate in opposite y-directions. Scale bar 10*µ*m.

**Video 3**: Movie of the actomyosin cortex during the third cell division. The cortex is marked by NMY-2::GFP. The P_1_ cell division does not exhibit chiral counter-rotating flows. The nature of the flow is similar to the one observed in case of P_0_ cell division. Scale bar 10*µ*m.

**Video 4**: Movie of the actomyosin cortex during the ABa and ABp cell divisions. The cortex is marked by NMY-2::GFP. ABa and Abp exhibit chiral counter-rotating flows during division similar to the flows observed in the AB cell division. Scale bar 10*µ*m.

**Video 5**: Movie of the actomyosin cortex during the EMS division. The cortex is marked by NMY-2::GFP. No chiral counter-rotating flows are observed during the EMS division. Scale bar 10*µ*m.

**Video 6**: Chiral counter-rotating cortical flows during the second and third cell division in *par-2; chin-1; lgl-1(RNAi)* embryos. The cortex is marked by NMY-2::GFP. Scale bar 10*µ*m.

**Video 7**: Cortical flows during the second and third cell division in control embryos. The second cell division exhibits chiral counter-rotating flows whereas the third cell division shows absence of chiral counter-rotating flows. Scale bar 10*µ*m.

**Video 8**: Cortical flows during the second and third cell division in *par-6 (RNAi)* embryos. No counter-rotating flows are observed during either of these cell divisions. Scale bar 10*µ*m.

**Video 9**: Movie of maximum intensity projection of dividing embryo from one-cell stage to 24 cell stage. Spindle apparatus and spindle poles are marked by TBB-2::mCherry(red) and membrane(grey) is marked by PH::GFP. Scale bar 10*µ*m.

**Video 10**: Cortical flows during the second cell division in *ect-2 (RNAi)* embryos. 4 hrs of *ect-2 (RNAi)* leads to reduction in chiral counter-rotating flow velocity (NMY-2::GFP, cortex marked in green and TBB-2::mCherry, spindle poles marked in red) Scale bar 10*µ*m.

**Video 11**: Cortical flows during the second cell division in *L4440* embryos (feeding control) (NMY-2::GFP, cortex marked in green and TBB-2::mCherry, spindle poles marked in red) Scale bar 10*µ*m.

**Video 12**: Cortical flows during the second cell division in *rga-3 (RNAi)* embryos. 6 hrs of *rga-3 (RNAi)* leads to increase in chiral counter-rotating flow velocity (NMY-2::GFP, cortex marked in green and TBB-2::mCherry, spindle poles marked in red). Scale bar 10*µ*m.

**Video 13**: Spindle skew in the AB cell imaged in control embryos (SWG63) by visualizing TBB-2::mCherry at restrictive temperature of 25°C. Scale bar 10*µ*m.

**Video 14**: Spindle skew in the AB cell imaged in the mutant *nmy-2(ts)* embryos (SWG204) by visualizing TBB-2::mCherry at restrictive temperature of 25°C. Scale bar 10*µ*m.

